# Reconnecting food production and consumption through redesigning food systems to support healthy diets

**DOI:** 10.1101/2025.11.25.690428

**Authors:** Caroline De Clerck, Tom Desmarez, Mathieu Delandmeter, Paulo César de Faccio Carvalho, Benjamin Dumont, Jérôme Bindelle

## Abstract

Balancing the social and environmental costs of food production with the needs of future populations in the face of climate change is the greatest challenge that agriculture is facing in the 21st century. The modification of eating habits towards more environmentally friendly and healthy diets is a key lever to meet this challenge. In 2019, the EAT-Lancet commission defined a universal guideline diet that should allow 9 billion people across the globe to eat healthily while respecting planetary boundaries.

In an attempt to reconnect cropping systems to such diets, we developed an innovative approach and designed a decision-making model to assist in the planning and optimization of cropping systems. This model evaluates their ability to supply a specific food system in accordance with the diet recommended by the EAT-Lancet Commission, while comparing vegan, ovo-lacto vegetarian, and omnivorous diets and minimizing the import and export of commodities.

Results show that longer and more diverse crop rotations are more likely to comply with the EAT-Lancet dietary requirements, especially if grazing animals are integrated within rotations. Integrated crop-livestock systems including temporary pasture and forage cover crops minimize the excess and deficit of all commodities (food and feed supplies), while reaching the required amount of daily calories and meeting the requirements in each food category. Crop rotations typically need to include rapeseed to provide oil for human consumption and oilseed meals for livestock, as well as a legume crop for pulses and several cereals. The model allows testing a wider range of dietary recommendations or assessing the impact of specific agro-ecological practices. It supports the design of multi-objective crop rotations aligned with dietary guidelines. By reconnecting food production and consumption following a healthy diet, the model provides practical solutions to sustain all three pillars of sustainability for future food systems.

## 1. Introduction

A variety of food systems evolved through History in response to local socio-environmental conditions. However, over the past decades, people with very different culinary traditions and cultural backgrounds stemming from their different food systems started eating increasingly similar diets (Khoury et al., 2014). This was encouraged by the increase in global incomes and wealth in many countries, which economically strengthened a middle-class and its demands for a more Westernized diet including more animal-based products and requiring more resources to produce (Godfray, 2015; Popkin, 1998).

This trend was accompanied by the evolution of production systems towards what is classified by Colonna et al. (2013)(Colonna et al., 2013) as the « agro-industrial » model. While the agro-industrial model was able to increase food production dramatically over the past century, it showed strong environmental limitations (Gerber et al., 2013). The agricultural sector has reached the point where land expansion and technological innovations have allowed the highest food energy production in History. However, this achievement has been met by an increasing specialization at the farm and landscape levels (Lemaire et al., 2014). The uniformity of these systems, and their reliance on large input-oriented chemical fertilizers, pesticides, and preventive use of antibiotics, systematically led to negative outcomes and vulnerabilities. Consequently, agriculture has become a cause of global environmental degradation (Ramankutty et al., 2018), leading current transgression of several planetary boundaries, within which mankind should remain in order to continue thriving for generations to come (Rockström et al., 2009). Over 48% of the food we consume today is grown under conditions that violate at least one of these planetary boundaries (Gerten et al., 2020). This calls for the need to redesign our food production systems and consumption patterns, inherently supported by the redesign of cropping systems.

The change of eating habits towards more environmental-friendly and healthy diets is a key lever to build a sustainable future (Gerten et al., 2020). While the acceptable total amount of animal-based food in the diets is still debated (Astrup et al., 2020), there is a large consensus that Westernized diets include excess meat and few fiber-rich plant-based foods from both the health and the environmental perspective (e.g. (Aleksandrowicz et al., 2016; Chai et al., 2019; Wellesley et al., 2015)). Diets containing high amounts of refined sugar, refined fats, oil and meat are known to greatly increase the incidence of type II diabetes, coronary heart diseases and other diseases that lower global life expectancies (Becerra-Tomás et al., 2020; Feskens et al., 2013; Tilman and Clark, 2014). Various expert- and modelling-based studies have come out in recent years to suggest how to redesign eating patterns. Stehfest et al. (Stehfest et al., 2009), aiming to reach important improvement in land use, greenhouse gases (GHG) emission and human health, studied the possibility of a global transition towards a diet containing less meat or even a complete switch to plant-based protein foods. Analyzing Mediterranean, vegetarian, pescetarian diets – three diets that are usually considered as interesting alternatives to a global-average diet - Tilman & Clark (Tilman and Clark, 2014) showed that all three alternative diets could reduce environmental impacts, particularly through lower GHG emissions. Alexander et al. (Alexander et al., 2016) defined an index for the Human Appropriation of Land for Food in order to specifically determine the effect of diets on agricultural land areas. They determined that the types of food commodities are more important than the quantity of food that is consumed in the determination of agricultural land use, largely because of the highest requirements for animal products. Following their findings, 55% less agricultural land would be needed if the world was to adopt the average Indian diet, with low meat intakes.

Willett and his collaborators of the EAT-Lancet Commission (Willett et al., 2019) took all these studies in consideration when proposing a universal guideline diet that would allow 9 billion people across the globe to eat healthily while respecting the planet boundaries. This diet mostly consists of fruits, vegetables and nuts that are usually supplied by horticulture and perennial agriculture. But also, of a large amount of products that rely on annual crop agriculture, such as whole grains, legumes, starchy roots and tubers, unsaturated oils, sugar. Their diet was designed to include no or low quantities of animal products (dairy products, meat, eggs, seafood). According to Willet et al. (Willett et al., 2019), to achieve its goals of sustainability, changes of eating pattern toward the diet they suggest must be considered in a context of overall change in consumption patterns and in citizens’ behavior and culture, such as large reduction of food losses and waste, and major improvements in food production practices.

In order to provide locally such healthy and sustainable diets, it appears urgent to redesign food and agricultural systems and one of the crucial steps is to reconnect what sustainable crop rotation can offer to what people should ideally eat. One of the key reasons is the risks associated with supply chains in a globalized market. This has been evident in regions such as North Africa and the Middle East during the Ukraine war, where disruptions to food imports occurred (Abay et al., 2023), and in the European Union during the COVID-19 pandemic, which exposed vulnerabilities in the supply of essential goods like medical supplies (Bown, 2022). These kinds of disruptions, particularly in the agricultural sector and food supply, are likely to intensify in the future due to climate change, threatening global food security (Gregory et al., 2005). Therefore, the model aims i.a. at minimizing excesses (possible exports) and deficits (required imports) of food and feed commodities.

Crop redistribution studies, investigating the allocation of crop species and activities to areas in agricultural landscapes (Memmah et al., 2015), involve complex decision-making processes that must balance multiple, often conflicting objectives to reduce trade-offs (Kaim et al., 2018). Several objectives can be prioritized in these strategies, including maximizing farmer profitability (Capitanescu et al., 2017; Galán-Martín et al., 2015), promoting circular agricultural systems (Van Zanten et al., 2023), enhancing water management practices (Femeena et al., 2018; Rulli et al., 2024; Singh, 2012), ensuring food security (Femeena et al., 2018; Van Zanten et al., 2023; Wang et al., 2022), and increasing agricultural productivity for both food and biofuel production (Femeena et al., 2018; Galán-Martín et al., 2017). These objectives often coexist within a framework of complex constraints, such as climate change adaptation (Klein et al., 2013), reducing environmental impacts (Capitanescu et al., 2017; Femeena et al., 2018), adherence to agricultural policies (Galán-Martín et al., 2015), and respecting crop rotation requirements (Galán-Martín et al., 2015). The integration of these factors is essential for developing effective, sustainable agricultural strategies that balance profitability with ecological stewardship.

Despite the potential benefits, crop redistribution and land optimization strategies are still underutilized, especially in more developed regions. Current research and applied models primarily focus on optimizing food supply, often without fully addressing other essential aspects of agricultural sustainability, such as promoting food autonomy (Dai et al., 2023; Kuzmanovski et al., 2019; Sali et al., 2016; Van Zanten et al., 2023). Moreover, only a limited number of studies explore the competition for land between human food production and livestock or other animal uses (Van Kernebeek et al., 2016; Van Zanten et al., 2023; Wang et al., 2024). Additionally, few models incorporate guidelines for healthy, sustainable diets into their frameworks (Rulli et al., 2024; Van Zanten et al., 2023; Wang et al., 2024), which could play a significant role in shaping more sustainable and nutritionally balanced agricultural landscapes.

In this study, we propose an innovative approach to optimize the design of cropping systems. Based on a participatory approach with experts from multiple disciplines, including farmers, we co-designed 40 diverse crop rotations according to regional climatic constraints, contrasting by their duration and crop diversity. Then, given these rotations as inputs, our decisional model allows to determine the use of crop (by-)products (*i*) to maximize the amount of dietary energy to satisfy human requirements, (*ii*) while minimizing the excesses and deficits in the different food and feed commodities, under the constraints (*iii*) of satisfying the eating patterns proposed by the EAT-Lancet commission which would allow the globe to eat healthily while respecting planet boundaries (Willett et al., 2019), (*iv*) in the contrasting contexts of an omnivorous, an ovo-lacto vegetarian and a vegan diets.

We use this conceptual model in the Hesbaye region of Belgium, characterized by (*i*) a current food system dominated by an agro-industrial model (UNDP, 2020) ; (*ii*) a consequent high index of environmental pressure on the planet; (*iii*) a densely populated situation; and (*iv*) a high human population index (human development index of 0.931 and 374 inhabitants per km^2^). This is a situation that we assumed as typical of food systems from developed countries with lack of long-term sustainability in addition to a disconnection between local food production and consumption.

The objectives were (*i*) to evaluate the capacity of our innovative decisional model to evaluate how diversified crop rotations can fulfill specific nutritional requirements while minimizing imports and exports of food and feed commodities; (*ii*) to determine how these criteria are impacted by the duration and diversity of crop rotations, including crop-livestock integration; and (*iii*) to analyze the effects of contrasting human diets on the optimal utilization of crop (by-)products.

## 2. Material and methods

### 2.1. Agronomic context

The agronomic context of the silty region of Hesbaye (Belgium) was considered. The region is characterized by a temperate Cfb climate according to the Köppen–Geiger classification (Peel et al., 2007). Soils are Cutanic Luvisol (Schad et al., 2015), silty, with favorable natural drainage.

### 2.2. Co-designing diverse crop rotations for targeted diets

We compared 40 crop rotations (Table 1) varying in duration (3 to 8 years) and in major crop species and types (corn for silage or grain, hard or soft winter or spring wheat, winter or spring barley, sugar beet, potatoes, rapeseed, spring or winter peas, oats, hemp, temporary grasslands).

**Table 1.** List of the 40 rotations co-designed by the experts for this study

| Code | Rotation # | Year 1 | Year 2 | Year 3 | Year 4 | Year 5 | Year 6 | Year 7 | Year 8 |
| --- | --- | --- | --- | --- | --- | --- | --- | --- | --- |
| 3.1 | 1 | Silage corn | Winter wheat | Winter Barley | - | - | - | - | - |
| 3.2 | 2 | Cereal corn | Winter wheat | Winter Barley | - | - | - | - | - |
| 3.3 | 3 | Sugar beet | Silage corn | Winter wheat | - | - | - | - | - |
| 3.4 | 4 | Sugar beet | Cereal corn | Winter wheat | - | - | - | - | - |
| 3.5 | 5 | Sugar beet | Winter wheat | Winter Barley | - | - | - | - | - |
| 3.6 | 6 | Potatoes | Winter wheat | Winter Barley | - | - | - | - | - |
| 3.7 | 7 | Rapeseed | Winter wheat | Winter Barley | - | - | - | - | - |
| 4.1 | 8 | Sugar beet | Winter wheat | Silage corn | Winter wheat | - | - | - | - |
| 4.2 | 9 | Sugar beet | Winter wheat | Cereal corn | Winter wheat | - | - | - | - |
| 4.3 | 10 | Potatoes | Winter wheat | Silage corn | Winter wheat | - | - | - | - |
| 4.4 | 11 | Potatoes | Winter wheat | Cereal corn | Winter wheat | - | - | - | - |
| 4.5 | 12 | Sugar beet | Winter wheat | Potatoes | Winter wheat | - | - | - | - |
| 4.6 | 13 | Sugar beet | Winter wheat | Rapeseed | Winter wheat | - | - | - | - |
| 4.7 | 14 | Potatoes | Winter wheat | Rapeseed | Winter wheat | - | - | - | - |
| 5.1 | 15 | Sugar beet | Winter wheat | Spring Pea | Winter wheat | Winter barley | - | - | - |
| 5.2 | 16 | Potatoes | Winter wheat | Spring Pea | Winter wheat | Winter barley | - | - | - |
| 5.3 | 17 | Rapeseed | Winter wheat | Spring Pea | Winter wheat | Winter barley | - | - | - |
| 6.1 | 18 | Sugar beet | Winter wheat | Rapeseed | Winter wheat | Silage corn | Winter barley | - | - |
| 6.2 | 19 | Sugar beet | Winter wheat | Rapeseed | Winter wheat | Cereal corn | Winter barley | - | - |
| 6.3 | 20 | Potatoes | Winter wheat | Rapeseed | Winter wheat | Silage corn | Winter barley | - | - |
| 6.4 | 21 | Potatoes | Winter wheat | Rapeseed | Winter wheat | Cereal corn | Winter barley | - | - |
| 6.5 | 22 | Sugar beet | Winter wheat | Rapeseed | Spring potatoes | Winter wheat | Winter barley | - | - |
| 6.6 | 23 | Potatoes | Winter wheat | Rapeseed | Winter pea | Silage corn | Winter barley | - | - |
| 6.7 | 24 | Potatoes | Winter wheat | Rapeseed | Winter pea | Cereal corn | Winter barley | - | - |
| 6.8 | 25 | Potatoes | Winter wheat | Rapeseed | W.wheat + W.pea | Silage corn | Winter barley | - | - |
| 6.9 | 26 | Potatoes | Winter wheat | Rapeseed | W.wheat + W.pea | Cereal corn | Winter barley | - | - |
| 6.10 | 27 | Sugar beet | W.wheat + W.pea | Rapeseed | Spring potatoes | Winter wheat | Winter barley | - | - |
| 7.1 | 28 | Sugar beet | Winter wheat | Rapeseed | Winter wheat | Sugar beet | Winter wheat | Winter barley | - |
| 7.2 | 29 | Potatoes | Winter wheat | Rapeseed | Winter wheat | Potatoes | Winter wheat | Winter barley | - |
| 7.3 | 30 | Sugar beet | Winter wheat | Rapeseed | W.wheat + W.pea | Sugar beet | Winter wheat | Winter barley | - |
| 7.4 | 31 | Sugar beet | Winter wheat | Rapeseed | W.wheat + W.pea | Potatoes | Winter wheat | Winter barley | - |
| 7.5 | 32 | Sugar beet | Winter wheat | Rapeseed | Winter wheat | Spring Peas | Winter wheat | Winter barley | - |
| 8.1 | 33 | Grassland<br>Temp. | Grassland Temp. | Silage corn | Winter wheat | Sugar beet | Winter wheat | Rapeseed | Winter wheat |
| 8.2 | 34 | Grassland<br>Temp. | Grassland Temp. | Cereal corn | Winter wheat | Sugar beet | Winter wheat | Rapeseed | Winter wheat |
| 8.3 | 35 | Grassland<br>Temp. | Grassland Temp. | Silage corn | Winter wheat | Potatoes | Winter wheat | Rapeseed | Winter wheat |
| 8.4 | 36 | Grassland<br>Temp. | Grassland Temp. | Cereal corn | Winter wheat | Potatoe | Winter wheat | Rapeseed | Winter wheat |
| 8.5 | 37 | Grassland<br>Temp. | Grassland Temp. | Cereal corn | Winter wheat | Faba bean | Winter wheat | Rapeseed | W.wheat +<br>W.pea |
| 8.6 | 38 | Camelina | Winter wheat | Faba beans | Winter wheat | Rapeseed | Winter wheat | Spring Pea | Oats |
| 8.7 | 39 | Camelina | Winter barley | Faba beans | Winter wheat | Rapeseed | Winter wheat | Spring Pea | Oats |
| 8.8 | 40 | Sugar beet | Winter wheat | Potatoes | Winter wheat | Rapeseed | W.wheat + W.pea | Silage corn | Winter wheat |

These crop rotations were designed in a participatory approach with agronomy, nutrition and ecology experts. The objective was to compare contrasting crop rotations that were all consistent from an agronomic perspective. Some of these rotations are commonly practiced by farmers following a conventional model in the loamy region of Hesbaye, while others are more innovative, featuring longer durations and a greater diversity of crops as often suggested in organic agriculture and agroecology. In order to maintain ecosystem services such as biodiversity, soil quality, nutrient management, water-holding capacity, control of weeds, diseases, and pests (Kremen and Miles, 2012; Lin, 2011), the following designing constraints were applied: (*i*) maximization of the inclusion of cover crops, (*ii*) a periodicity of legumes greater than 3 years, (*iii*) and the alternance of botanical families (i.e. *Solanaceae*, *Brassicaceae*, *Amaranthaceae*, *Fabaceae*), except for *Poaceae*.

No constraint was set either on fertility or soil management. The productivity of each crop species and type was computed from the yield statistics in Hesbaye between 2014 and 2018 (Statbel, 2024). From the yield of each main commodity, by-products production levels were also derived (e.g. straw from cereal grain or oilseed cake from rapeseed) to yield the total commodities produced per crop species and types, including cover crops and temporary grasslands.

### 2.3. Optimization process to match targeted diets

An innovative approach was drawn to optimize the design of the crop rotations, with the aim of satisfying the requirement of specific diets. The use of a given commodity as food or feed was not decided *a priori*, but resulted from an optimization process described hereafter (Fig. 1).

**Figure 1.**
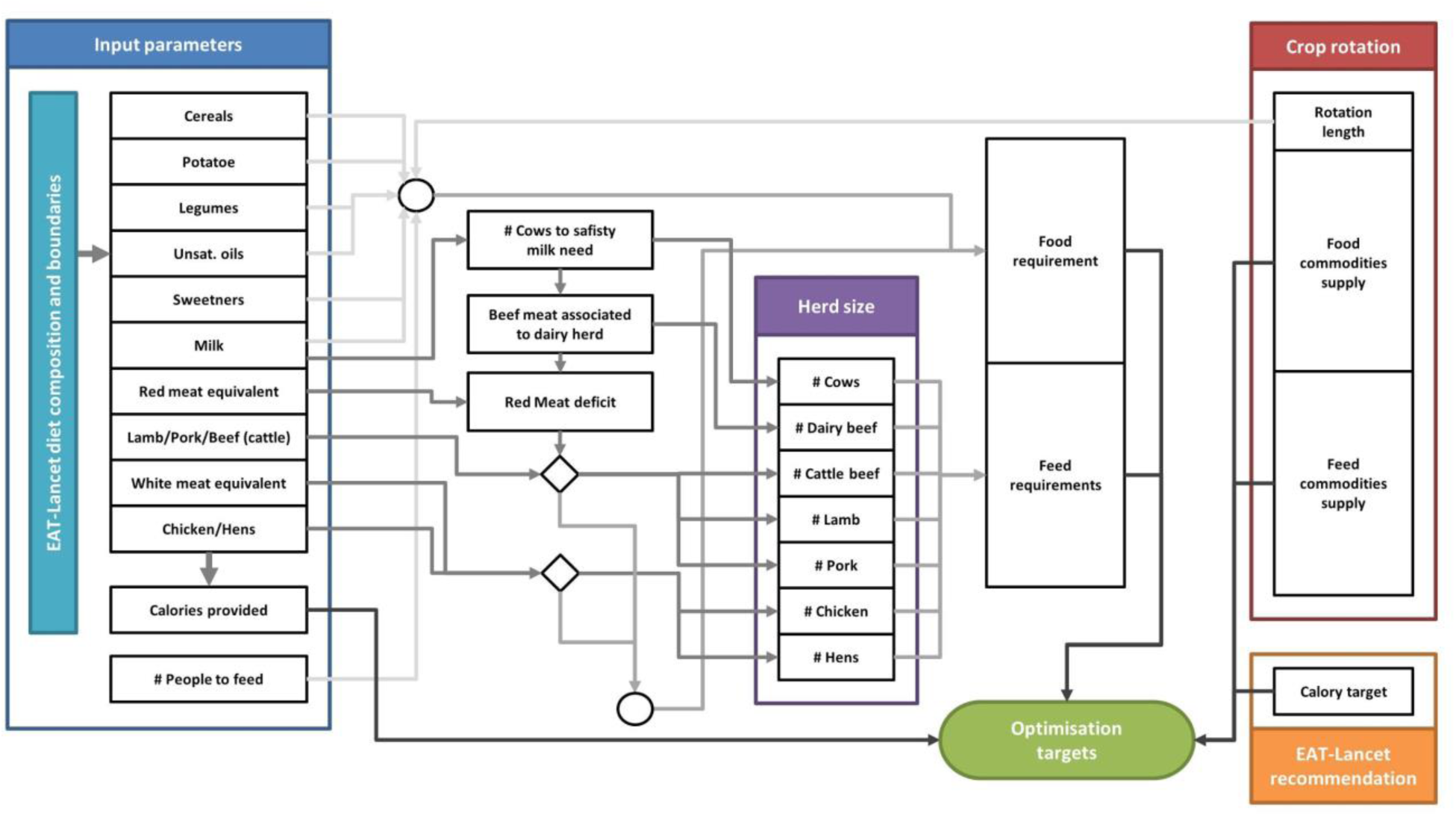
Optimization model to best allocate the different food category of the EAT-Lancet diet according to a given crop rotation, in order to minimize imports and exports.

Using the innovative crop rotations previously designed as input, the decisional model allows us to determine the use of crop (by-)products. It has two objectives: (*i*) maximizing the amount of dietary energy to satisfy human requirements, and (*ii*) minimizing the excesses and deficits in the different food and feed commodities per ha of land used. This optimization problem has two main constraints: (*i*) satisfying the eating patterns proposed by the EAT-Lancet commission, (*ii*) in the contrasting contexts of an omnivorous, an ovo-lacto vegetarian and a vegan diet. These objectives and constraints are detailed below.

By making use of what is produced by the crops in the rotation as food or feed, as well as the by-products, the main goal was to fulfill a dietary energy target (kcal) to satisfy human requirements. As recommended in Willett et al. (2019), a total amount of 2500 kcal day^-1^ was considered as the target achievable considering the agronomical productions at field scale. The other objective consisted in minimizing the excesses and deficits in the different food and feed commodities, including the one potentially required but not produced in the rotation.

A first constraint in the optimization process was to respect the specifications of the EAT Lancet diet (Willett et al., 2019) for each general food item category relevant in the context of field production (Table 2). The ranges defined were used as prior information (parameter *a priori* distribution) of our optimization process. Vegetables, fruits and tree nuts were considered as food commodities that had to be produced from outside the modelled crop rotations and fixed to the average values suggested in Willett et al. (2019) of 300, 200 and 25 g.d^-1^, providing 78, 126 and 149 kcal.d^-1^, respectively. As made possible in the same study, dairy fats were included in milk. Lard or tallow, fish, peanuts, soy foods and palm oil requirements were set to zero. The number of people being fed per ha was also a parameter, with an *a priori* distribution ranging from 1 to 100.

**Table 2.**
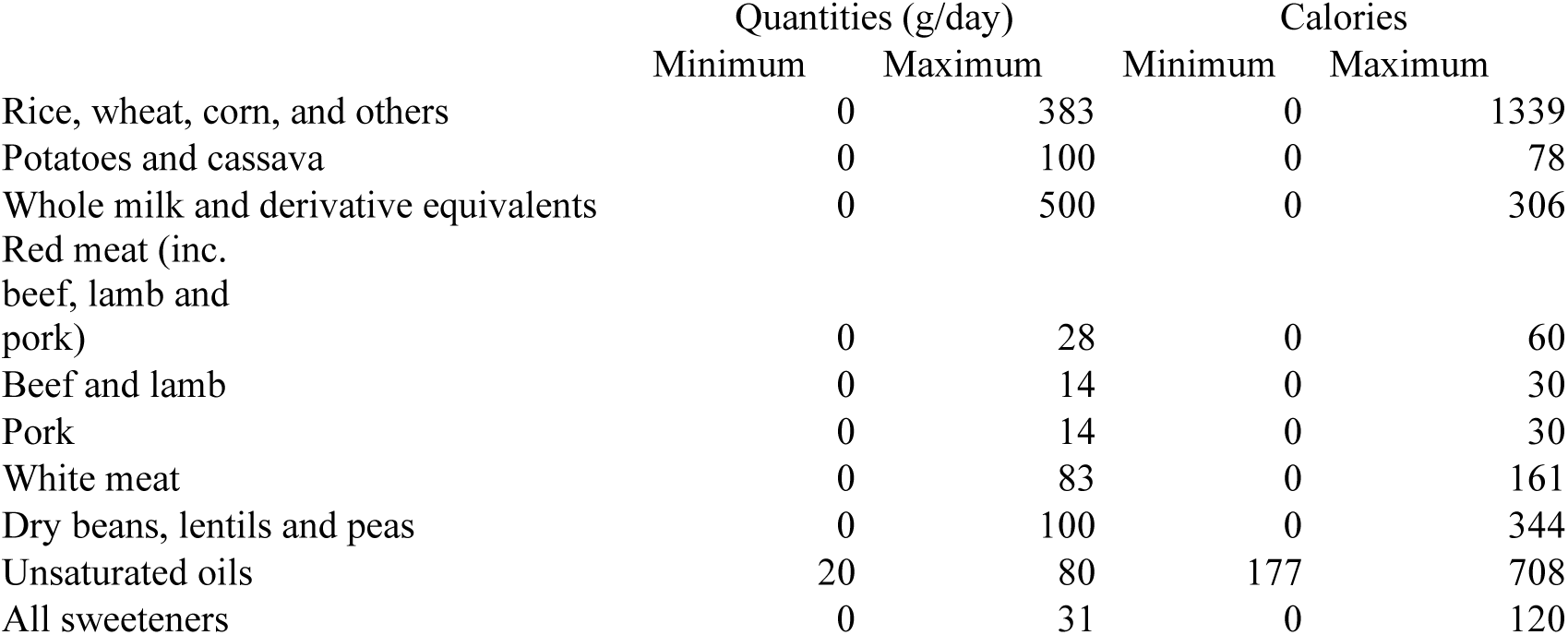
: EAT-Lancet diet considered in this study

A second constraint was to respect specific diets: an omnivorous, an ovo-lacto-vegetarian and a vegan diet. In each scenario, appropriate modifications were provided to the algorithms to consider livestock categories being fed or not and their respective products entering the balance of the diet.

The feed requirements for animal-based food production were calculated for the omnivorous diet, while this step was skipped for the vegan diet and adjusted for ovo-lacto vegetarian diets. The number of dairy cows needed to supply milk was estimated using life cycles, herd dynamics, and the annual production of milk, meat, and eggs per functional unit for each livestock species, as provided by Wilkinson (Wilkinson, 2011).

The resulting yields of red meat from dairy beef cattle and culled dairy cows were compared to the total red meat requirements. If necessary, additional red meat was produced from pork, lamb, and beef cattle. Similarly, the number of chickens and hens required to produce white meat (chicken meat and eggs) was calculated. Outputs from animal by-products, such as meat from culled breeding females, were included for cattle, sheep, and pigs, but not for poultry. Annual requirements for grassland forage, human-edible crops, and crop by-products to feed the resulting herds were calculated based on forage and concentrate requirements and composition data (Wilkinson, 2011). The human-edible crops used for animal feed were subtracted from the plant-based food commodities available for human consumption, enabling the final calculation of the optimal diet based on this specific feed/food ratio. This process was iteratively repeated during the optimization, over several cycles, until an optimal solution was found.

The algorithms were programmed in MatLab (MatLab *version 7.5.0 (R2007b)*. Natick, Massachusetts: The MathWorks Inc.). The optimization process was performed using the Differential Evolution Adaptive Metropolis (DREAM) algorithm (Vrugt et al., 2009). Detailed descriptions of the DREAM algorithm have been published in (Vrugt, 2016; Vrugt et al., 2009). DREAM has been successfully used in a wide range of scientific disciplines (Vrugt, 2016) among which in agronomy and crop modelling (Duchene et al., 2021; Dumont et al., 2014). The functions to compute the use of crop rotation to match a diet model can be obtained by contacting the authors. The DREAM source codes were obtained from the developer (Jasper A. Vrugt, personal communication).

### 2.4. Assessment of optimized crop productions and diet

The resulting diets matching the crop rotations best under the constraint of the omnivorous, ovo-lacto vegetarian or vegan option of Willet et al (Willett et al., 2019) specifications were assessed using several criteria provided as outputs from the model: (*i*) the number of people supported per ha per year, (*ii*) the amount of calories produced by the crops used for feed or food purposes, (*iii*) the amount of calories required from outside the system, (*iv*) the size of the animal herds supported by the rotation expressed in livestock units (Eurostat, 2024). The nutritional value of the diet was assessed by means of the amount (g or mg. d^-1^) and share (%) of animal-sourced protein, iron and zinc in the diets. Animal-sourced protein is commonly considered to address the question of sustainability of eating habits owing to the higher environmental impact of livestock farming. It is also important because of the role of animal-based foods in providing an adequate balance of bioavailable essential amino acids. Animal-sourced iron and zinc allows to address the bioavailability of various minerals in which plant-based foods are usually deficient from a nutrition perspective. Finally, the different components of the Nutri-Score and the Nutri-Score itself (ranging from A to E) of each resulting diets was calculated (Julia and Hercberg, 2017): energy density (kJ/100g), sugars (g/100g), saturated fatty acids (g/100g), Na (mg/100 g), fruits, vegetables, pulses, nuts rapeseed, walnut and olive oil (%), fiber (g/100g), total protein (g/100g). The latter were calculated using the USDA National Nutrient Database (USDA, 2024). Food commodities that were not to be provided by the crop rotations under any crop rotation and optimization result (dark green, red orange and other vegetables, tree nuts and fruits) were set to similar levels for all crop rotations by providing systematically the required amounts suggested by Willet et al. (2019). Their nutrient content was also taken from the USDA National Nutrient Database. The architecture of the model can be found in Figure 1.

A PCA analysis was conducted using the singular value decomposition approach. It was implemented under the R language (R Core Team, 2023). The *prcomp* function from the "*factoextra*" library was used. Data were centered and standardized as recommended. Optimized diet composition (including elements composing red and white meat), excess/deficit in commodities, the nutritional value of food (iron, protein, zinc, etc.), the different components of the Nutri-Score and the number of people fed constituted the analyzed data, to understand the variability in the data set. Diet type, share of crops in each rotation and herd compositions (expressed in LSU) were used to structure the dataset, as parsing arguments, to further understand how the different variables were related.

## 3. Results and discussion

### 3.1. Land use

Results show that short-term rotations are less self-sufficient and lead to higher commodity excesses or deficits (Fig. 2, S1, S2). Three-years rotations display particular large deficits in different food and feed commodities in all the diets considered (depending on the crops that were cultivated). Long-term rotations very importantly reduce the deficits, even if results strongly differ between rotations of the same duration with different crop species (Fig. 2, S1, S2).

**Figure 2.**
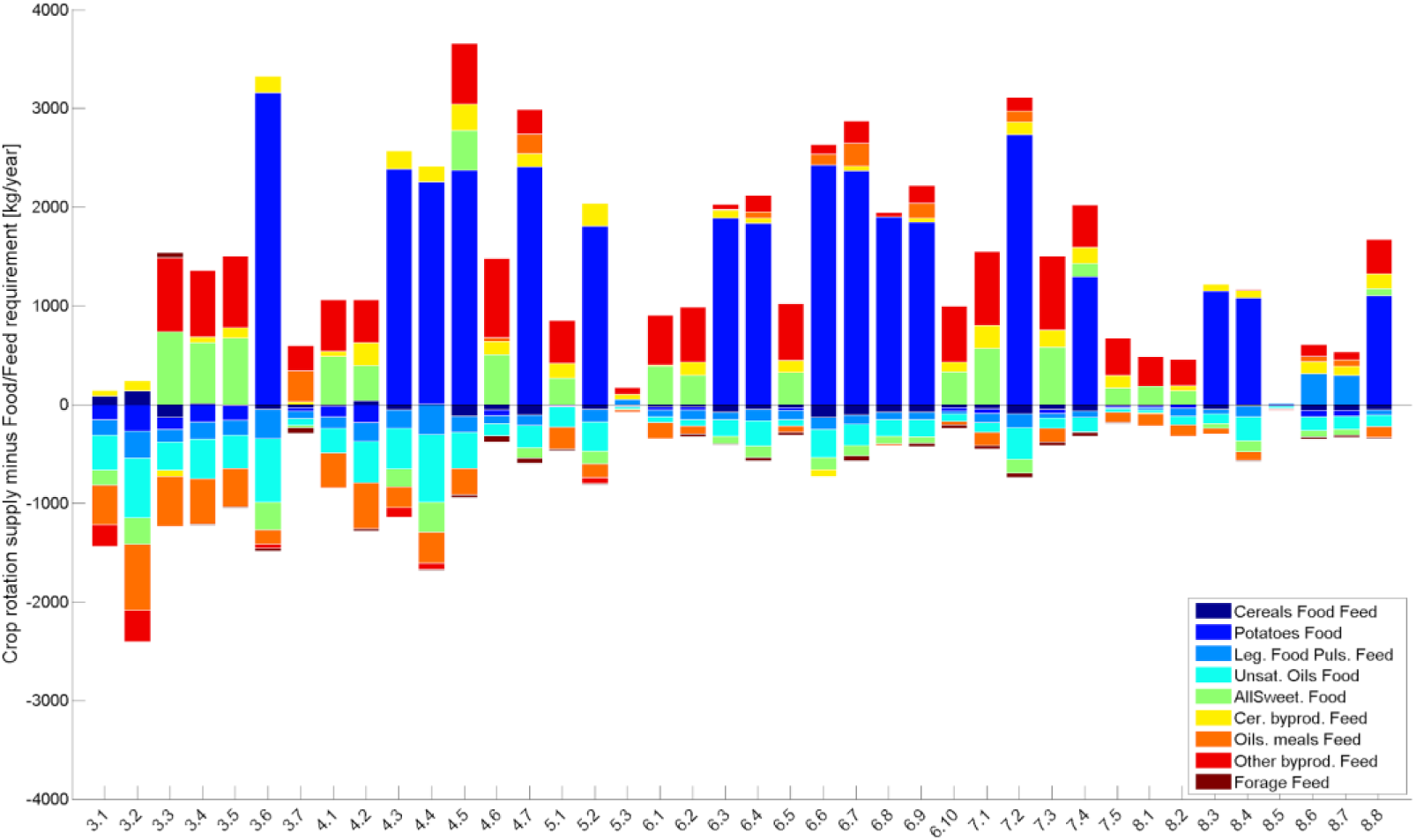
Excesses and deficits in food and feed commodities compared to the EAT Lancet requirements, for all rotations considering an omnivorous diet. In rotations labels, first numbers indicate duration in years.

Following our model results, the ideal rotation must contain rapeseed to supply oil for humans and oilseed meals for livestock (for the omnivorous and ovo-lacto vegetarian diets), one legume crop for pulses and several cereals as main energy source in the diets designed according to Willet et al. (2019) (Fig. 2, S1, S2).

For all diets and in all tested crop rotations, cereals are almost at the equilibrium (Fig. 2, S1, S2). Our model was built to ensure that when there are no potatoes in the rotation, it minimizes the amount of tubers required in the diet so that deficits are as low as possible. Yet potatoes are always an issue when they are included in the rotations as they always lead to overproduction (on average 1989 ± 585 kg/ha/year). Unsaturated oils are always in deficit. Concerning sweeteners, they are in excess when sugar beet is cropped, but in deficit when this crop is absent of the rotation (Fig. 2, S1, S2). In the omnivorous and ovo-lacto vegetarian diets, oilseed meals are in deficit when no rapeseed is included in the rotations or when rapeseed is present but legumes crops are absent because the frequency of rapeseed is too low in these crops rotations (Fig. 2, S1).

Vegan diets can feed more people on average (32.7 ± 9.7 versus 25.7 ± 6.2 for Omnivorous and 29.6 ± 6.5 for ovo-lacto vegetarian diets) (Fig. S3, S4 and S5), and deficits are globally lower than for the two diets allowing the inclusion of animal-based foods (Fig. S2). Yet some commodities are in large excess, while many others have to be supplied from outside (Fig. S2). Excesses are generally higher for all rotations with vegan diets than with diets including animal-based foods, and increasing rotation duration does not permit to decrease these excesses. They are particularly important for crop by-products that are usually used as feeds: forage (2441.9 ± 1774.3 kg/ha/year for Vegan, 4.2 ± 99.7 kg/ha/year for ovo-lacto vegetarian and - 18.5 ± 20.7 kg/ha/year for Omnivorous) oilseed meals (322.6 ± 252.4 kg/ha/year for Vegan, -150.7 ± 301.2 kg/ha/year for ovo-lacto vegetarian and -111.5 ± 212.6 kg/ha/year for Omnivorous), brans (301.6 ± 53.2 kg/ha/year for Vegan, 138.7 ± 111.8 kg/ha/year for ovo-lacto vegetarian, 99.2 ± 77.8 kg/ha/year for Omnivorous), etc. (Fig. S2). Those are strong clues indicating that the vegan diets would not exhibit the most efficient land use. Nevertheless, from a farming system perspective, the share of biomass produced in excess under the constraint of vegan diets could be used as green manure or as input for agro-sourced fuel production.

These observations, showing that diets with small amounts of animal proteins are more efficient in terms of land use, are consistent with the results of other studies (Peters et al., 2016; Van Kernebeek et al., 2016; Van Zanten et al., 2018). However, for ovo-lacto vegetarian diets, the question of the fate of animals used for eggs and milk production fully remains (e.g. 46.5 ± 32.3 kg/ha/year of dairy meat). In the omnivorous diets, their terminal destination as meat at the end of their productive life allows for a higher efficiency in the use of dairy cattle raising questions about vegetarian diets in terms of food system efficiency and circularity (Van Zanten et al., 2019).

The results highlight the importance of livestock and principally ruminants, to convert inedible byproducts of the human food industry and forage into edible milk, eggs and meat (Eisler et al., 2014; Van Kernebeek et al., 2016).

### 3.2. Herd composition

In the 40 tested rotations, the share of milk and beef in omnivorous and milk in ovo-lacto vegetarian diets is correlated to the presence of temporary grasslands but also to corn silage, forage cover crops and crops providing fiber-rich byproducts (Fig. 5, S6 and S7).

Under the omnivorous scenario, ruminants are not always the most adapted species to be integrated with crops in a perspective of feed vs. food competition (Fig. 3). Pork and chicken are proposed by the optimization process when cereal crops and co-products for feed are abundant in the rotation. Regarding the chicken-to-egg ratio for poultry, the model is often falling for chicken, as it seems to be more efficient to use the feed energy captured by the system than eggs. This is probably due to the short life cycle of chicken considered here (42 days) and the use of males and females alike, while for laying hens, only newly hatched females are kept and it takes almost 5 months for pullets to start laying their first egg (Fig. S8-S9).

**Figure 3.**
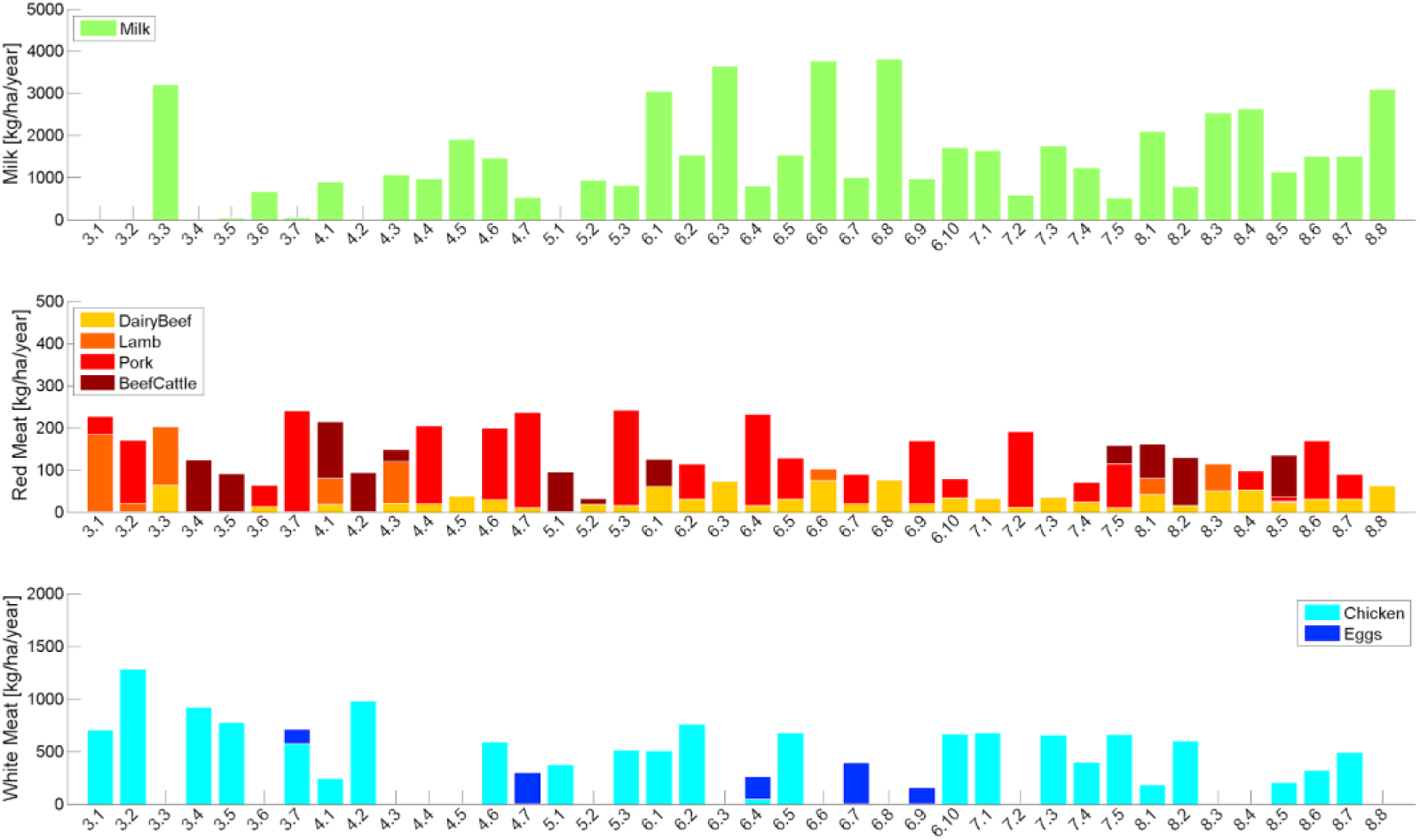
Quantities of animal products (milk, red meat, white meat or eggs) in kg/ha/year produced for each crop rotation in the case of an omnivorous diet complying with the recommendations of the EAT-Lancet commission

Interestingly, we observe that not all crop rotations in the omnivorous diet are to produce milk. Even if meat is produced and consumed in all rotations, the sources of red and white meat differ (Fig. 3). In the case of the ovo-lacto vegetarian diet, milk is produced in all rotations, but eggs are not. When no eggs are produced, the production of milk is usually more important (Fig. S10).

It is commonly accepted that land use for the production of a unit of protein is generally lower with plants than animal sources (Bai et al., 2021; De Vries and De Boer, 2010; Nijdam et al., 2012) but it is strongly linked to the considered livestock species and livestock production system. Dairy foods have the most efficient feed to food protein conversion ratio, followed by eggs and chicken then pork and lamb. Beef production is the least efficient way of supplying animal proteins through animal feeding in current farming systems (Smil, 2002). However, ruminants can take advantage of forage, grazing lands and food processing residues that are not digestible for non-ruminant species, converting these so-called low opportunity-cost feeds, farm animals recycle biomass and nutrients into the food system that would otherwise be lost to food production (Van Zanten et al., 2019), expressing better feed efficiency than non-ruminants when food/feed competition is eliminated (Van Zanten et al., 2016). The integration of temporary grasslands and forage cover crops in rotation will thus allow a higher production of beef at a lower expense.

In addition, these temporary grasslands will add some biodiversity in the rotation and the agricultural landscape. They can also have fertilizing and weed control effects, trough the presence of legumes and animal manure and through competition for light and grazing, respectively (Schuster et al., 2019). They are also highly effective in restoring soil health, reducing soil erosion and under appropriate management result in more C sequestration than emissions (Crème et al., 2020). Incorporating forages and ruminants into agroecosystems can also improve soil ecological function, through the minimization of tillage, inorganic fertilizers and biocides, and enhance wildlife habitat (Teague et al., 2016).

### 3.3. Energy intake

Excesses in dietary energy were never encountered in the optimized diets for all rotations but the model minimizes the deficits (Fig. 4, S11 and S12). In all rotations, the commodities produced were indeed able to provide at least 97.5, 97.2 and 95.7% of the recommended energy intake set to 2054 kcal per day for the omnivorous, ovo-lacto vegetarian and vegan diets respectively. As a reminder, the total supply in energy in the EAT-Lancet diet is 2500 kcal day^-1^ with, in our model, tree nuts, vegetables and fruits providing the additional energy from outside the cropping system.

**Figure 4.**
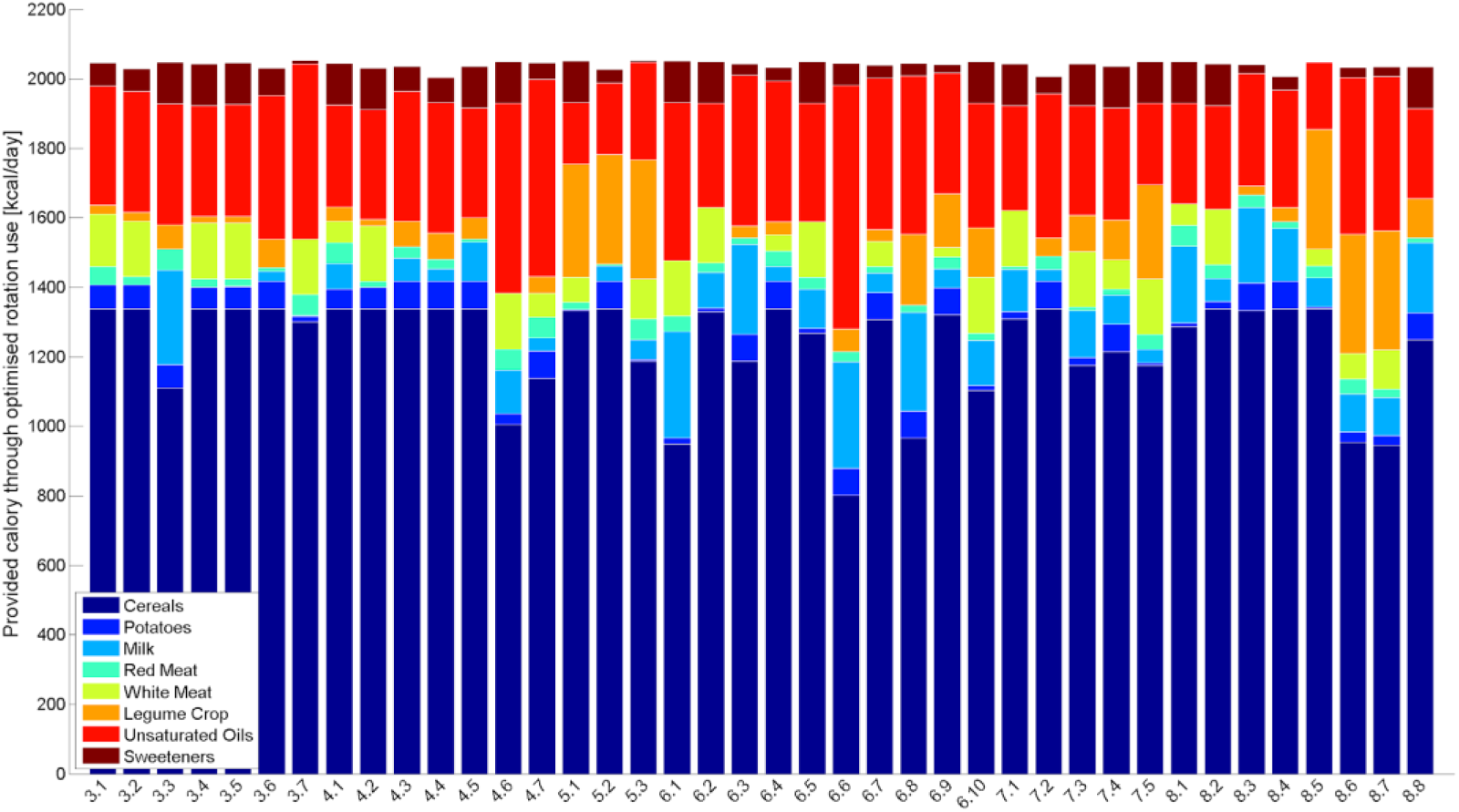
Share of the different food commodities in the provision of energy by all the rotations in the case of an omnivorous diet

In all crop rotations and for all diets, the production of cereals provides the largest share of energy, followed by unsaturated oils. When potatoes are included in the rotation, their dietary use is maximized with still overproduction as discussed above (Fig. 2, S1 and S2). Under a vegan scenario, the model always maximizes the share of cereals in the diets, up to the upper threshold defined by the EAT-Lancet diet, to compensate for the absence of energy-dense animal-source foods (Figure 4).

### 3.4. Nutritional profile

From the optimized use of farming systems production as feed and food, nutritional profiles were derived (see supplementary material), allowing for an integrating analysis from an agricultural and nutritive perspective (Fig. 5, S6 and S7).

**Figure 5.**
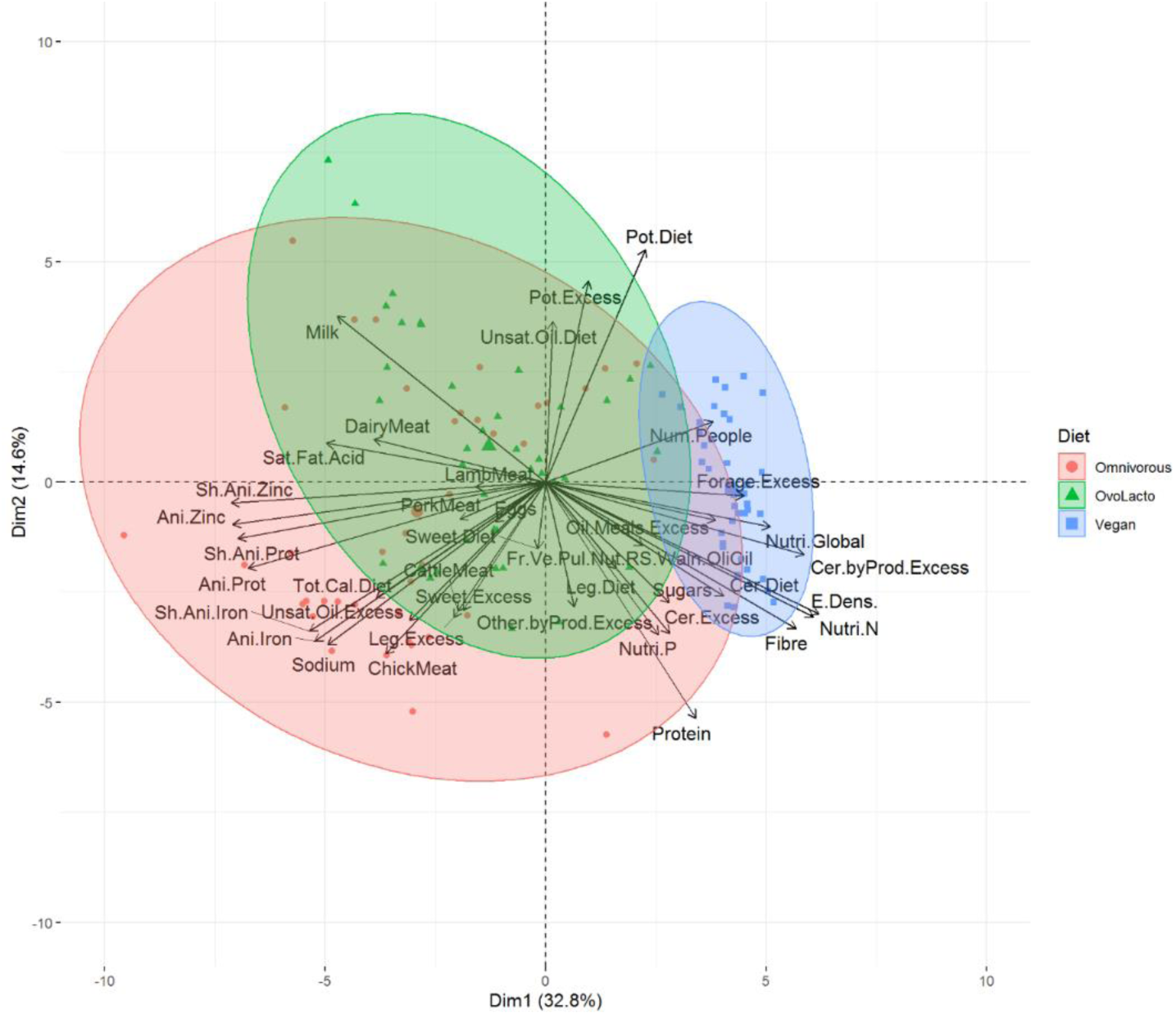
PCA analysis of crop rotations according to the matching with the requirements of the EAT-Lancet commission for omnivorous, ovo-lacto vegetarian and vegan diets. Clusters highlight the type of diet. All abbreviations are described in Table 1 (in sup mat.)

All crop rotations scored A for their Nutri-Score whatever the optimized diet (omnivorous, ovo-lacto vegetarian, vegan), indicating high nutritional quality and lower risk of developing health issues linked to nutrition (Deschasaux et al., 2020; Mozaffarian et al., 2021). This is in accordance with the objective of the EAT Lancet commission, aiming to reduce the risk of diet-related obesity and other non-communicable diseases, including coronary heart disease, stroke, and diabetes.

The share of animal proteins in diets was also correlated with herd sizes. Contents in animal protein were almost always higher in omnivorous diets, except when more pork and lamb than beef are produced as source of red meat (Fig. S14).

However, some rotations applied to the ovo-lacto vegetarian diets, but also to the omnivorous diets, had values that were really low in terms of total protein supply (with minima of 54.5 g/day for Omnivorous and 54.2 g/day for ovo-lacto vegetarian).These values are borderline according to the WHO recommendations (WHO et al., 2007) of 58 g of total protein per day for 70-kg individuals, especially if less than ⅓ is supplied as animal-based protein for their higher biological value. This is unfortunately the case of most low protein scoring diets, and the case of all vegan diets, obviously (with a minimum of 40.6 g of proteins/day). Interestingly, eight rotations supply over ⅓ of animal-based protein in the omnivorous diets while still complying with the EAT-Lancet recommendations. Van Zanten et al. (2018) have shown that the best land use in terms of number of people supported is reached when people eat 20-25 g animal protein/day. In our study, some farming systems are a bit higher (eg. crop rotations 3.7 is providing 28.7 g/day of animal proteins in the omnivorous diet and 29.8 g/day of animal proteins in the ovo-lacto vegetarian diet). The least efficient at this level (crop rotation 8.5) provided 13.9 and 10.5 g animal proteins per day for the omnivorous and ovo-lacto vegetarian diets, respectively. Several studies have shown that higher plant protein intake was associated with lower mortality (Budhathoki et al., 2019; Song et al., 2016). However, attention must be paid to the provision of minerals, micronutrients and vitamins. In addition to protein, iron (Fe) is one of the most critical nutritional requirements to be met in most diets in humans. Content in Fe were similar in all diets, with the exceptions of ovo-lacto vegetarian diets favoring high consumption levels of eggs (Fig. S13), which are among the richest in Fe of the animal-based foods (USDA, 2024).

It is interesting to consider as it could be an indicator of the level of mineral elements of animal origin. Its deficiency is more often the result of low bioavailability than low total supply (Hambræus, 1999). Iron and some other minerals are more bioavailable in animal products. Plant-based foods can be rich in Fe, but are usually deficient in bioavailable forms of iron from a nutrition perspective, especially for adult women. As shown by Päivärinta et al., (Päivärinta et al., 2020) flexitarian diets containing a small proportion of animal products would provide healthy and more sustainable alternatives for current, mostly animal-based diets.

### 3.5. Consequences to current specialized production systems in Belgium

Interestingly, the last rotation assessed (rotation 8.8) is virtually the most representative of the soil occupation in Hesbaye - the percentage of each crop present in the rotation mimics the actual land use by major crops in Belgium - although this soil occupation is actually achieved with a combination of several types of 3, 4 or 5-year rotations.

Presented results clearly show that the present-day land use in Belgium has to be improved in order to support a more sustainable diet. In particular, there is a large excess in the production of potatoes which is exported, when the country presents a deficit of animal feeds which are compensated through imports.

Improvements should also concern livestock production. The herd sizes proposed by the optimized model are far from the herd sizes that are usually encountered in the Hesbaye (Riera et al., 2019). For rotation 8.8, the optimized herd size would be 0.48 cows and 0.32 dairy beefs and would allow to feed 25.7 people per hectare. This would correspond to an average of 61.8 g of red meat intake per day. Herd sizes proposed in our optimally used rotation would follow the same trend than the suggestion made by Riera et al. (Riera et al., 2019), showing that herd size in Belgium could be reduced to provide the animal protein needed.

Under the vegan diet constraint, conclusions are more difficult to draw for the Hesbaye case study, due to the large excesses in feed crops or non-edible crop parts that cannot be exploited through animal production. A rotation producing no forage (e.g. rotation 3.7) seems to be well adapted to vegan dietary patterns. However, as stated before, some of the commodities produced in excess could be used to supply agro-sourced fuel production (Hoogwijk et al., 2003). In this case, rotations 5.3 and 8.5 are also good options to consider for the vegan diet.

Considering all these exposed results, one rotation (rotation 8.5) seemed to be highly adapted to the specific Hesbaye context, minimizing deficits and excesses (food waste or exports) for all commodities, under the omnivorous and ovo-lacto vegetarian diets. Such a rotation is long (8 years), containing two years of temporary grasslands, cereal corn, three years of winter wheat in which one is associated with winter peas, faba beans and rapeseed. Typically, this rotation fits under the definition of integrated crop-livestock system.

## 4. Conclusions

A conceptual model allowing to assess the ability of crop rotations to answer precise dietary targets was designed in this study. Such a tool offers a practical solution to anticipate required changes in the agricultural systems. The design of innovative cropping systems is needed to allow future food systems to address all three pillars of sustainability altogether, while reconnecting healthy diets with sustainable agricultural systems.

Aside of tree nuts, vegetables and fruits which are still to be produced from outside the modelled crop rotations, we demonstrated *in silico* that it is possible to design crop rotations based on the EAT Lancet dietary requirements sustaining, in the most optimistic case, the whole population of Belgium with staple foods (cereals, pulses, oils and animal-based foods) produced locally. But only long-term field trials, potentially coupled with the use of soil-crop models, will determine the resistance and resilience of the systems from an agronomical perspective.

This modeling work also shows the value of including animals in the farming system for their ability to complete the nutrient cycle, recycle co-products and supply micronutrients essential to the proper functioning of the human body.

Yet, all these results are to be nuanced as there are many debates about the relevancy of some nutritional recommendations proposed by the EAT Lancet Commission diet and the fact that it is very severe in terms of energy intakes and animal based foods leading to a potential decline in the supply of micronutrients such as iron, zinc and vitamins B_6_ and B_12_ (Beal et al., 2023). In parallel to the changes required regarding diets, major efforts are needed to improve the sustainability of food-production practices, and cropping systems have to be considered from an environmental point of view, to reduce pesticide use and the waste from production and consumption (Tuomisto, 2019).

Longer and more diversified rotations - alternating spring and winter crops, crops with different root systems and nutritional needs, or dense and hoed crops - are inherently more sustainable from an agroecological perspective, as they allow to naturally reduce pest and disease pressure and better exploit soil resources. Furthermore, the presence of cover crops within cropping is valuable at many levels, as they can be used as green manure or nitrate traps, as well as bringing soil protection against erosion or providing additional fodder capacity for example. While they might bring unused excesses of biomass under the vegan diet constraint, cover crops rich in *Fabaceae* allow to increase soil fertility and compensate for the lack of restitution of animal dejection.

Diet recommendations are universal, but are to be adapted to local conditions, as the characteristics of truly sustainable diets remains context-specific and depending on environmental and socio-economic factors (Milner and Green, 2018). In the present paper, the model was applied to design innovative crop rotations corresponding to the recommendations of the EAT Lancet commission, but it opens the door to broader uses and could be a powerful tool to determine the best choices to make when designing crop rotations adapted to socio-economic and/or environmental conditions.

## 5. Declarations

### 5.3. Authors contributions

BJ and DB conceptualized the study and designed the conceptual framework of the model; BJ and DC performed the data collection; DB implemented the model and conducted the statistical analysis; BJ, DB, DC, DM, DT and CPCF analyzed the data; BJ, DB, DC, DM, DT and CPCF produced the manuscript.

### 5.4. Availability of data and code

The raw data and code used in this study are available upon request. Interested researchers can contact the corresponding author via email to access these resources, which include data sets, calculation files, experimental protocols, and any other relevant materials.

