## Supplementary Material for "Reconnecting food production and consumption through redesigning food systems to support healthy diets"

### 1. Supplementary results

#### 1.1. General overview

Globally, results show that shorter crop rotations display larger deficits in different food and feed commodities, depending upon the crops that are cultivated (Figures 2, S1, S2). For omnivorous and ovo-lacto vegetarian diets, deficits and excesses tend to decrease with the duration and the diversity of crops in the rotation, showing that longer rotations tend to better address the recommendations to reduce animal-based food systems.

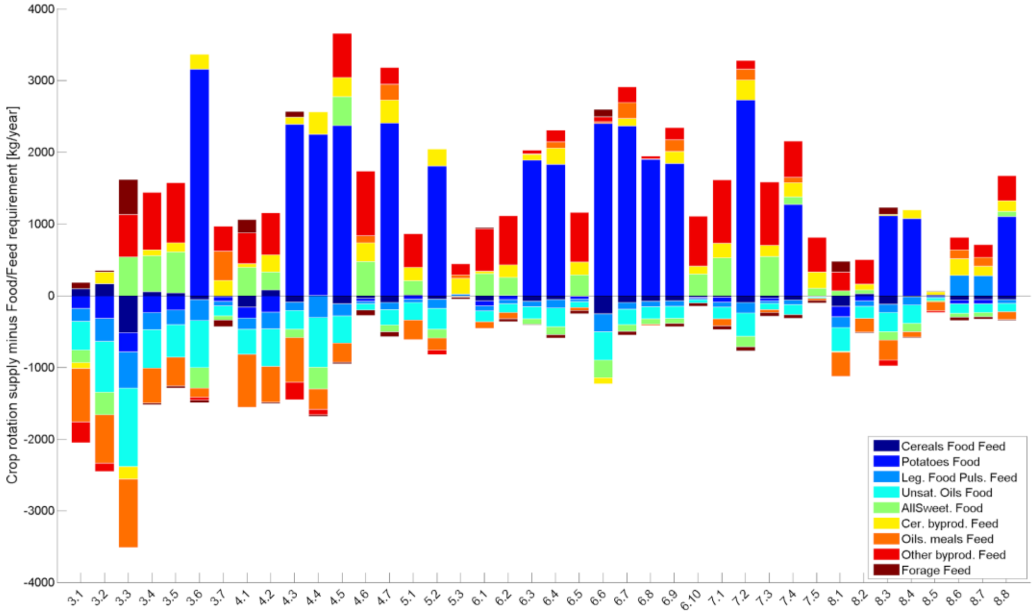


*Figure S1. Excesses and deficits in food and feed commodities compared to the EAT Lancet requirements, for all rotations considering an ovo-lacto vegetarian diet*

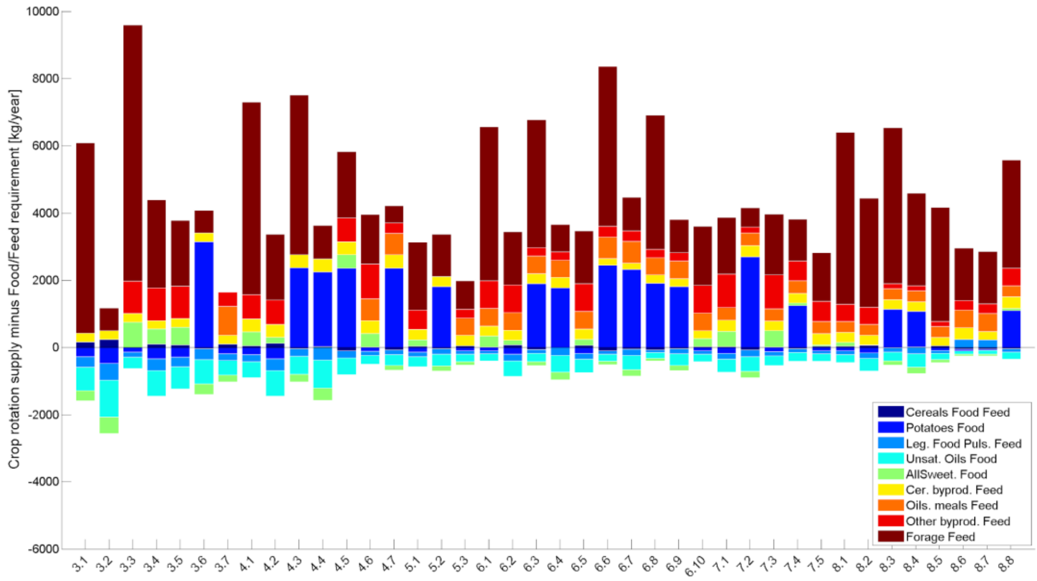


*Figure S2 Excesses and deficits in food and feed commodities compared to the EAT Lancet requirements, for all rotations considering a vegan diet*

In vegan diets, deficits are globally lower than for the two diets allowing the inclusion of animal-based foods. But the trend towards a reduction in excesses with increasing duration is less marked, notably due to the fact that excesses are generally associated to unused forage.

For all diets and in all rotations, cereals are almost at the equilibrium. Potatoes are systematically in excess when they are present. When there are no potatoes in the rotation, the model minimizes the amount of tubers required in the diet so that deficits are as low as possible. Unsaturated oils are almost always in deficit. All sweeteners are always in deficit when no sugar beet is cropped, and in excess when this crop is present once per rotation.  In terms of numbers of people supported per area, the global range spans from 18 to 65 people per ha across the different rotations and diets. The vegan diets allow on average to support higher numbers of people to be fed as highlighted on the PCA results (Figure 5). Moreover, deficits remain to be compensated are provided from outside of the rotation assessed (Figures S3, S4, S5).

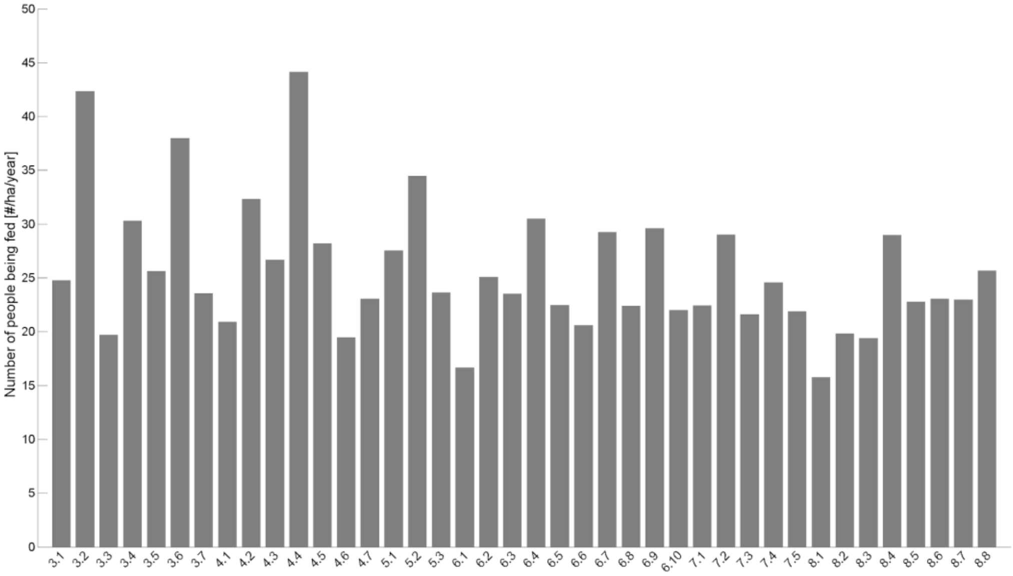


*Figure S.3. Number of people fed per ha per year according to the different crop rotations in the case of an omnivorous diet complying with the recommendations of the EAT-Lancet commission*

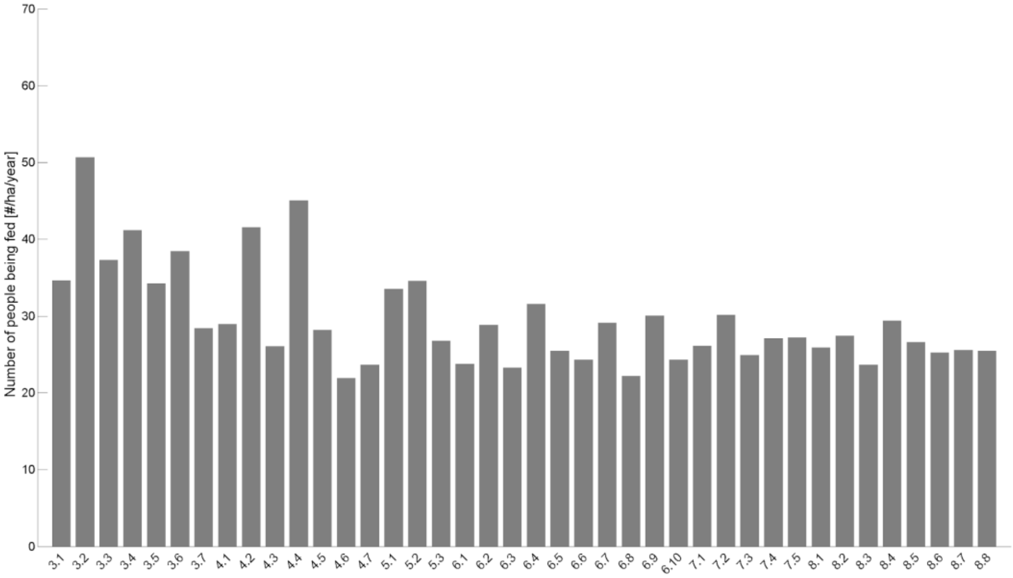


*Figure S.4. Number of people fed per ha per year according to the different crop rotations in the case of an ovo-lacto vegetarian diet complying with the recommendations of the EAT-Lancet commission*

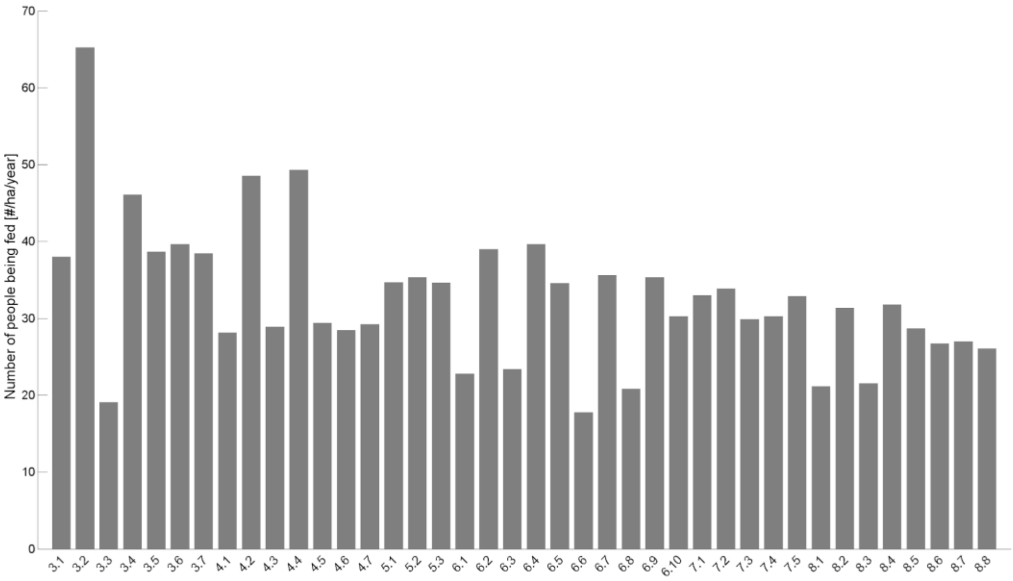


*Figure S.5. Number of people fed per ha per year according to the different crop rotations in the case of a vegan diet complying with the recommendations of the EAT-Lancet commission*

The omnivorous and ovo-lacto vegetarian diets are deficient in oilseed meals when no rapeseed is included in the rotations (rot. 3.1 to 3.6, 4.1 to 4.5, 5.1 and 5.2) or when rapeseed is present but legumes crops are absent (in rot. 4.7, 6.1, 6.2, 6.5, 7.1, 8.1 to 8.4). Rotations 7.3, 7.5 and 8.8. appear to be an exception to this rule.

### 1.2. Specificities observed under the different diets

Regarding animal-based foods, in the omnivorous diet, meat is produced and consumed in all rotations (Figure 3). Interestingly, not all rotations are to produce milk ((rot. 3.1, 3.2, 3.4, 4.2 and 5.1 (Figure 3).

In rotations 4.5, 6.3, 6.8 and 8.8, only milk is produced (no eggs) and dairy beef is the only source of meat. Production of eggs is chosen by the optimization model in rotations 3.7, 4.7, 6.4, 6.7, 6.9 in combination with chicken, and alone in rotations 4.7, 6.7 and 6.9. Chicken is produced in all rotations except rotations 3.3, 3.6, 4.4, 4.5, 5.2, 6.3, 6.6, 6.8, 7.2, 8.3, 8.4 and 8.8. The highest amounts of milk, chicken and red meats are produced respectively in rotations 3.2, 6.6 and 5.3. In rotations with temporary grasslands (8.1 to 8.5) herds tend to converge towards diets with a higher share of beef. The presence of corn silage and cover crops in the rotation seemed linked to the increase in milk and dairy beef (Figures S6 and S7, respectively).

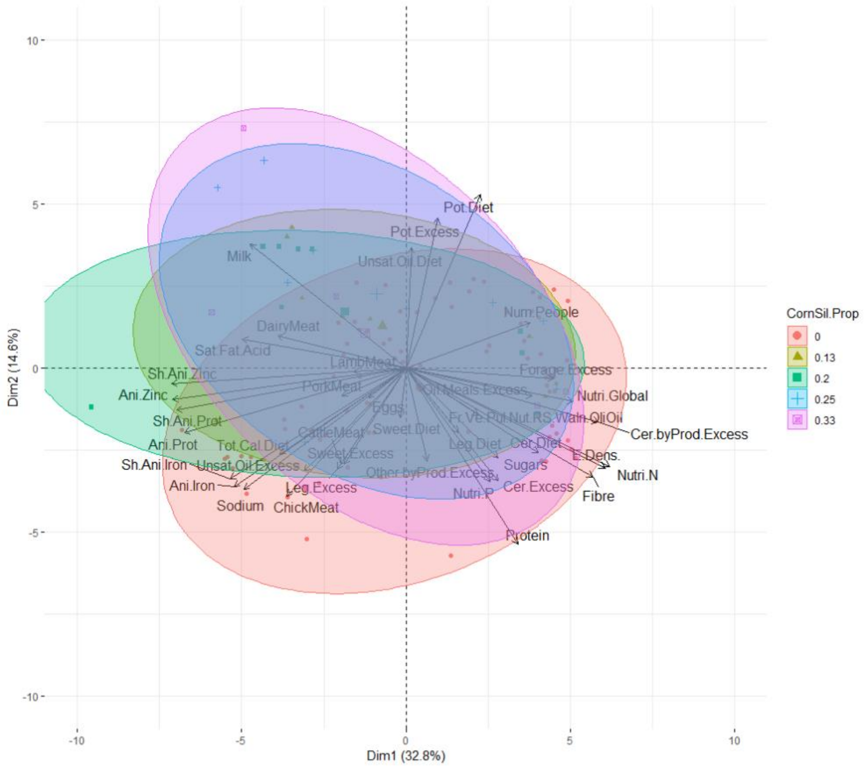


*Figure S6. PCA analysis of crop rotations according to the matching with the requirements of the EAT-Lancet commission for omnivorous, ovo-lacto vegetarian and vegan diets. Clusters highlight the share of corn silage as main crop in the rotations. All abbreviations are described in Table 1.*

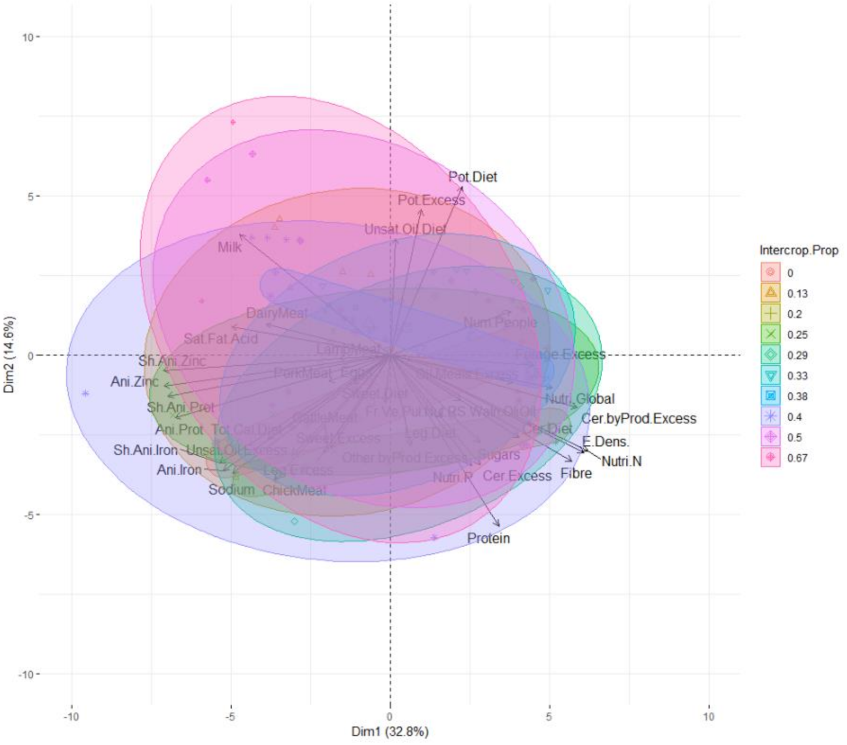


*Figure S7. PCA analysis of crop rotations according to the matching with the requirements of the EAT-Lancet commission for omnivorous, ovo-lacto vegetarian and vegan diets. Clusters highlight the frequency of cover crops in the rotations. All abbreviations are described in Table 1.*

*Table 1. List of abbreviations used in this study and associated measurement units*

| Abbreviation | Signification (units) |
|---|---|
| Tot.Cal.Diet | Total calories in the diet (kgCal/day) |
| Cer.Diet | amount of calory provided by cereals to the diet (kgCal/day) |
| Pot.Diet | amount of calory provided by potatoes to the diet (kgCal/day) |
| Leg.Diet | amount of calory provided by legumesto the diet (kgCal/day) |
| Unsat.Oil.Diet | amount of calory provided by unsaturated oils to the diet (kgCal/day) |
| Sweet.Diet | amount of calory provided by sweeterners to the diet (kgCal/day) |
| Milk | Quantity of Milk (kg/ha/year) |
| DairyMeat | Quantity of dairy meat (kg/ha/year) |
| LambMeat | Quantity of lamb meat (kg/ha/year) |
| PorkMeat | Quantity of pork meat (kg/ha/year) |
| CattleMeat | Quantity of cattle meat (kg/ha/year) |
| ChickMeat | Quantity of chicken meat (kg/ha/year) |
| Eggs | Quantity of eggs(kg/ha/year) |

| | |
|---|---|
| Cows | number of cows (LSU) |
| Dairy.Beef | number of dairy beefs (LSU) |
| Lamb | number of lambs (LSU) |
| Pork | number of pigs(LSU) |
| Cattle | number of cattles (LSU) |
| Chicken | number of chicken (LSU) |
| Hens | number of hens (LSU) |
| Cer.ExcDef | Excess or deficit in cereals (kg/ha/year) |
| Pot.ExcDef | Excess or deficit in potatoes (kg/ha/year) |
| Leg.ExcDef | Excess or deficit in legumes (kg/ha/year) |
| Unsat.Oil.ExcDef | Excess or deficit in unsaturated oils (kg/ha/year) |
| Sweet.ExcDef | Excess or deficit in sweetners(kg/ha/year) |
| Cer.byProd.ExcDef | Excess or deficit in cereal byproducts (kg/ha/year) |
| Oil.Meals.ExcDef | Excess or deficit in oil meals (kg/ha/year) |
| Other.byProd.ExcDef | Excess or deficit in other byproducts (kg/ha/year) |
| Forage.ExcDef | Excess or deficit in forage (kg/ha/year) |
| Cer.Excess | Cereal excess (kg/ha/year) |
| Pot.Excess | Potato excess (kg/ha/year) |
| Leg.Excess | Legume excess (kg/ha/year) |
| Unsat.Oil.Excess | Unsaturated oil excess (kg/ha/year) |
| Sweet.Excess | Sweetners excess (kg/ha/year) |
| Cer.byProd.Excess | Cereal byproducts excess (kg/ha/year) |
| Oil.Meals.Excess | Oil meals excess (kg/ha/year) |
| Other.byProd.Excess | Other byproduct excess (kg/ha/year) |
| Forage.Excess | Forage excess (kg/ha/year) |
| Sh.Ani.Prot | share of animal protein (g/g) |
| Ani.Prot | animal protein (g/d) |
| Sh.Ani.Iron | share of animal iron (mg/mg) |
| Ani.Iron | animal iron (mg/d) |
| Sh.Ani.Zinc | share of animal zinc (mg/mg) |
| Ani.Zinc | animal zinc (mg/d) |
| E.Dens. | energy density (kJ/100g) |
| Sugars | sugars (g/100g) |
| Sat.Fat.Acid | saturated fatty acids (g/100g) |
| Sodium | sodium (mg/100 g) |
| Nutri.N | nutriscore-N |
| Fr.Ve.Pul.Nut.RS.Waln.OliOil | Fruits, vegetables, pulses, nuts rapeseed, walnut and olive oil (%) |
| Fibre | Fibre (g/100g) |
| Protein | Protein (g/100g) |
| Nutri.P | Nutriscore-P |
| Nutri.Global | Global nutriscore |

Rotations 3.2, 3.6, 4.4, 5.2 are able to feed the highest number of people (Figure S3), again assuming that the deficits are provided from outside of the rotation. It is interesting to note that rotations 3.6 and 5.2, characterized by the lowest total herd sizes expressed in LSU, are not the rotations feeding the lowest number of people (Figures S8, S9).

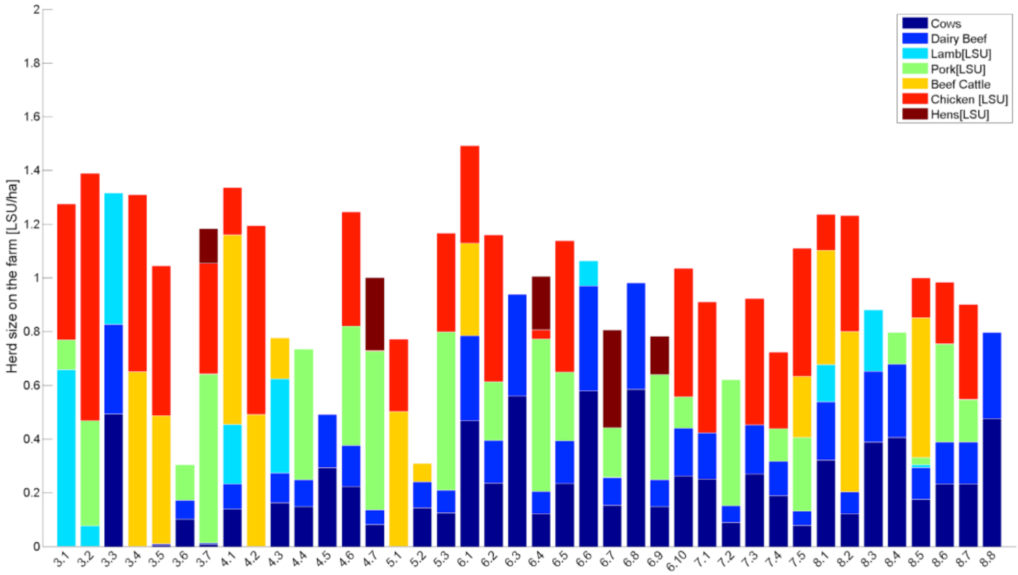


*Figure S.8. Herd composition for the different crop rotations in the case of an omnivorous diet complying with the recommendations of the EAT-Lancet commission (in LSU/ha)*

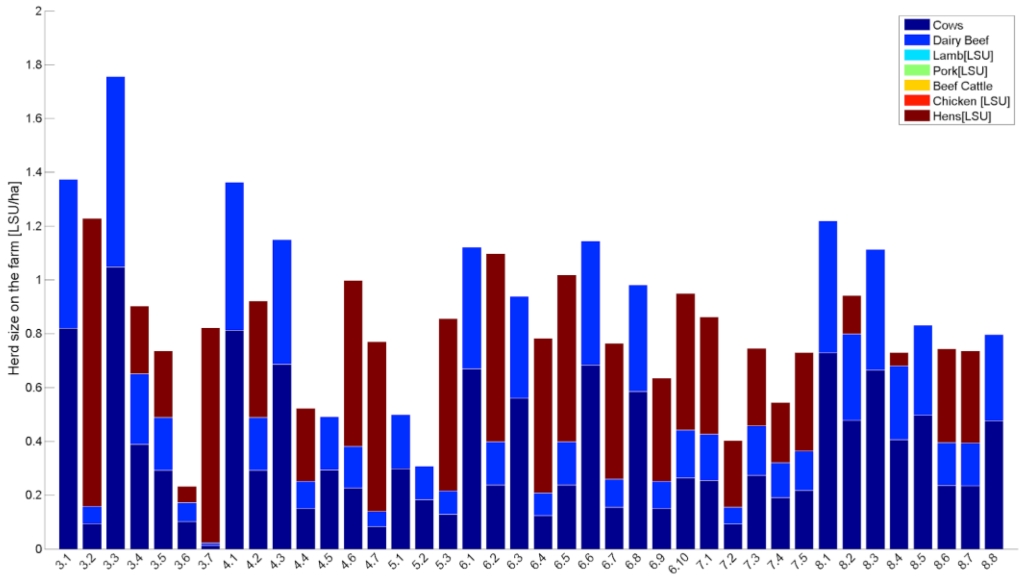


*Figure S.9. Herd composition for the different crops rotations in the case of an ovo-lacto vegetarian diet complying with the recommendations of the EAT-Lancet commission (in LSU/ha)*

In the ovo-lacto vegetarian diet, meat is produced in all rotations where dairy cattle is present but not consumed as dairy beef from culled cows, male calves and non-replacement heifers is a co-product of milk production (Figure S10).

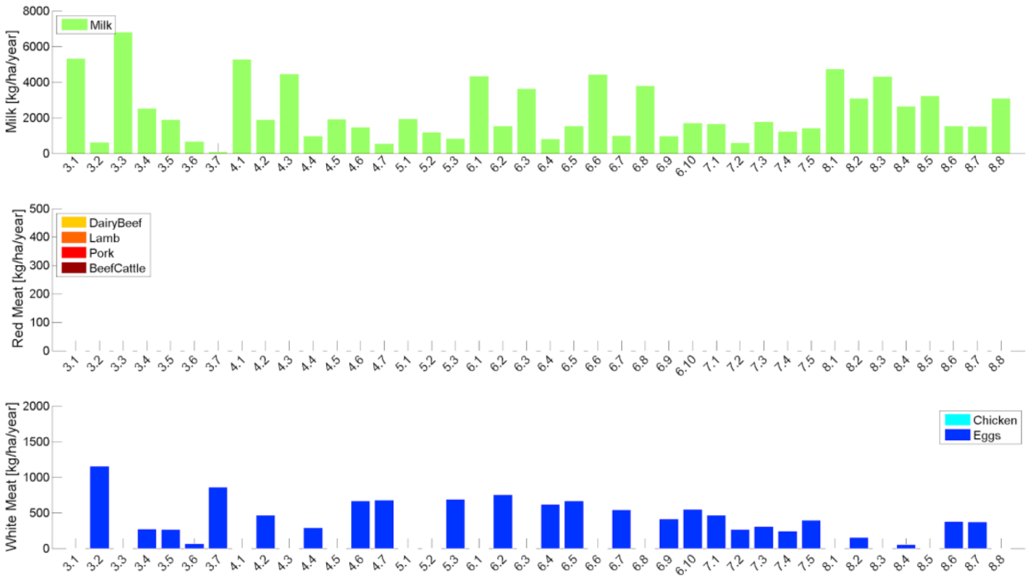


*Figure S10. Quantities of animal products (milk, red meat, white meat or eggs) in kg/ha/year produced for each crop rotation in the case of an ovo-lacto vegetarian diet complying with the recommendations of the EAT-Lancet commission*

Milk is produced in all rotations but eggs are not. They are produced in rotations 3.2, 3.4 to 3.7, 4.2, 4.4, 4.6, 4.7, 5.3, 6.2, 6.4, 6.5, 6.7, 6.9, 6.10, 7.1 to 7.5, 8.2, 8.4, 8.6 and 8.7. When no eggs are produced in the rotation, the production of milk is usually more important, except in rotation 4.5, 5.1 and 5.2. The greatest amounts of milk and eggs are produced in rotations 3.3 and 3.2. Rotation 3.2 is also the one feeding the highest number of people eating an ovo-lacto vegetarian diet; namely 50.7 people/ha. It is interesting to note that here again rotations 3.6 and 5.2 are characterized by the lowest herd size (0.23 and 0.3 LSU/ha, respectively) but are not the rotations feeding the lowest number of people (38.5 and 34.6 people/ha, respectively), even when compared to other rotations of the same duration in the case of 3-year rotations.

In the vegan diet, excesses are particularly important for crop by-products that are usually used as feeds: forage, oilseed meals, brans, etc. Not all crop products are indeed edible for humans. Rotation 3.7, producing no forage because it has neither intercrops, nor temporary grasslands is again an exception to this rule. Rotation 3.2 is, with 65.3 people/ha, the one feeding the highest number of people eating a vegan diet but also all diets combined (Figure S5).

### 1.3. Calory provision

Figures 4, S11 and S12 show that, for each rotation, the commodities produced are able to provide at least 97.5, 97.2 and 95.7% of the recommended energy intake set to 2054 kcal per day for the omnivorous, ovo-lacto vegetarian and vegan diets respectively.  As a reminder, this value represents the dietary energy that should be provided by the food commodities produced by the crops in the rotation only, thus tree nuts, fruits and vegetables set aside.

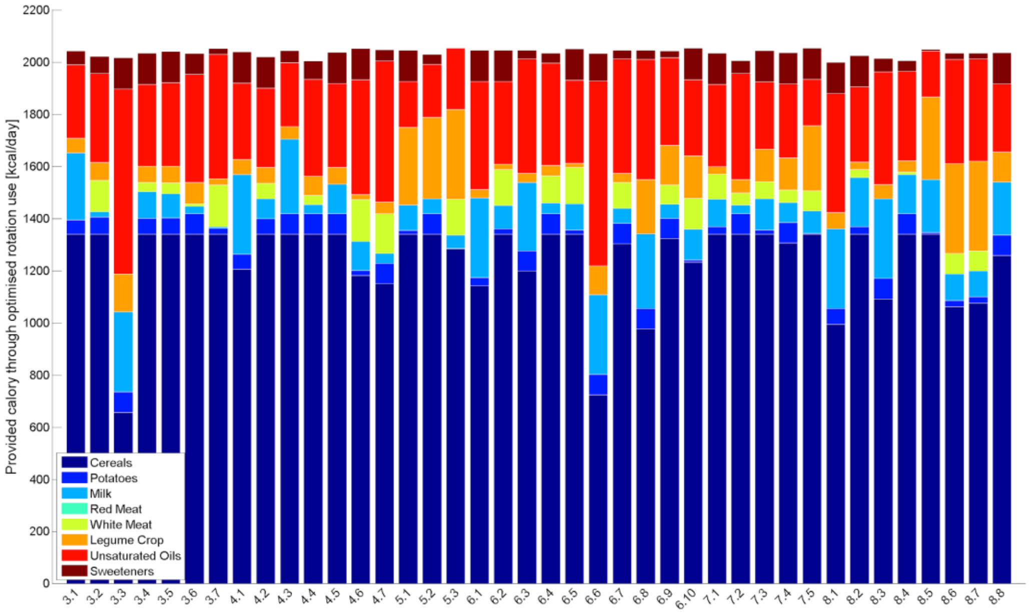


*Figure S11.* Share of the different food commodities in the provision of energy by all the rotations in the case of an ovo-lacto vegetarian diet

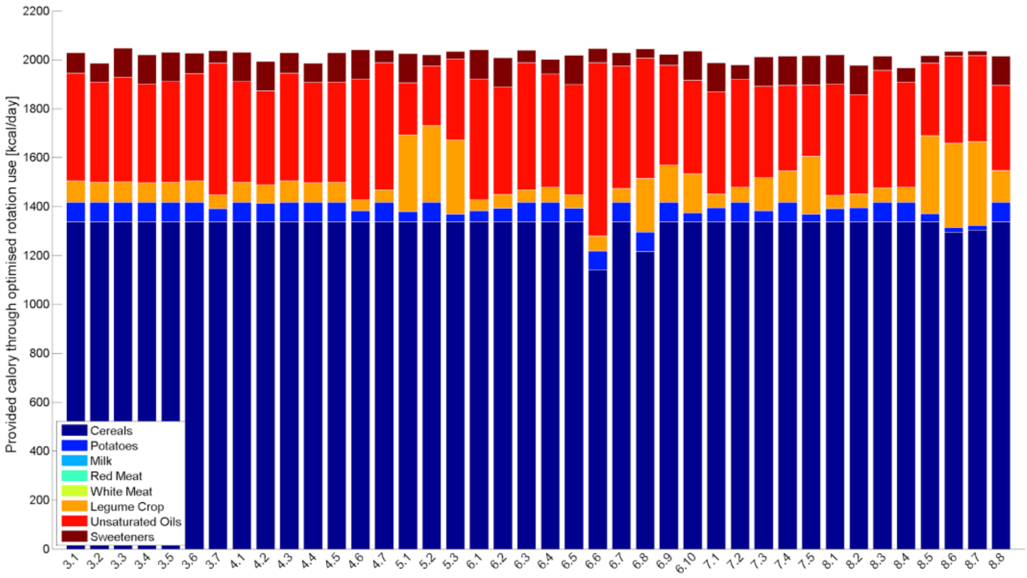


*Figure S12.* Share of the different food commodities in the provision of energy by all the rotation in the case of a vegan diet

In all rotations and for all diets, the production of cereals is optimized to provide the largest proportion of energy, followed by unsaturated oils. In some rotations (5.1, 5.2, 5.3, 7.5, 8.5, 8.6, 8.7), legume crops are also important sources of energy. Legume crops contribute obviously to a greater part of the calories intake for the vegan diet. In rotations without potatoes, the amount of energy having to be supplied by potatoes is set as a minimum by the model, as it will be otherwise considered as a deficit. This effect is particularly marked with the increase in the duration of the rotations.

Although limited in absolute value, the most important deficits are found in omnivorous diets based on rotations 4.4, 7.2 and 8.4, where potatoes are cropped and animal products contribute to a limited share of the diets. In omnivorous diets based on rotations 3.7, 5.1 and 5.3, producing no potatoes but using higher levels of meat, the deficit in energy is lower. In the ovo-lacto vegetarian diet, important deficits are also found for rotations 4.4, 7.2 and 8.4 but also for rotation 8.1. In the vegan diet, calories deficits are higher for rotations 3.2, 4.2, 4.4, 7.1, 7.2, 8.2 and 8.4.

### 1.4. Nutritional provision

Regarding their nutritional qualities, all rotations scored A for their Nutri-Score whatever the optimized diet (omnivorous, ovo-lacto, vegan). The negative components of the Nutri-Score (energy density, sugars, saturated fatty acids, Na), were more influenced by the rotations than the positive ones (protein, fiber, fruits, nuts and vegetables). The maximum value for the positive component of the Nutri-Score was 15 (rot. 3.1 vegan diet) due to its high use of unsaturated oils and pulses, but it scored at 10 for most rotations in all diets. The lowest positive score (7) was obtained for rotation 6.6 in the omnivorous diet because of its lower fiber (<4.7 g/100g) and protein density (<6.4 g/100g), providing neither eggs nor chicken, a large amount of milk and some lamb. The same trend was followed in the ovo-lacto vegetarian diet, where rotations provide no eggs but a large amount of milk (e.g rot. 3.3, 6.6 and 8.1) had smaller positive scores (7 to 8). The negative component of the Nutri-Score ranged between 4 (for rotations 3.3, 6.1, 6.3, 6.6 and 6.8 in the omnivorous diet and rotations 3.3, 4.1, 6.1, 6.3, 6.6, 6.8, 8.1 and 8.3 in the ovo-lacto vegetarian diet) and 8. All rotations scoring the highest for this negative component of the Nutri-Score did so when supplying vegan diets. The latter were actually impacted by their higher energy density stemming from a higher use of oils to fulfill the requirements.

In addition to the Nutri-Score, we looked at the iron and proteins supplied from animal-based foods in the omnivorous and ovo-lacto vegetarian diets (Figures S13 and S14). Content in Fe were almost always higher in omnivorous diets, except for the rotations 3.2, 3.6, 3.7, 4.4, 4.6, 4.7, 5.3, 6.2, 6.4, 6.5, 6.7, 6.9, 7.2 and 8.6, rich in eggs in the ovo-lacto vegetarian diet.

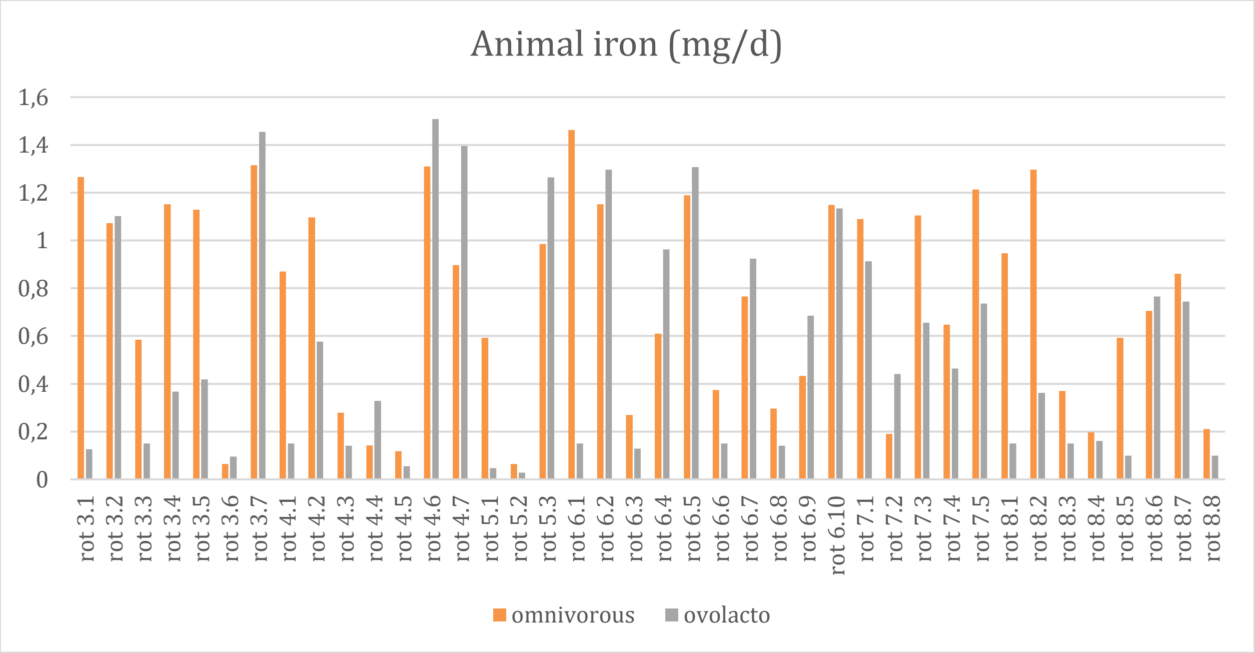


*Figure S.13. Amount of animal iron (mg/d) provided by the different crop rotations in the case of omnivorous (in orange) and ovo-lacto vegetarian (in gray) diets complying with the recommendations of the EAT-Lancet commission*

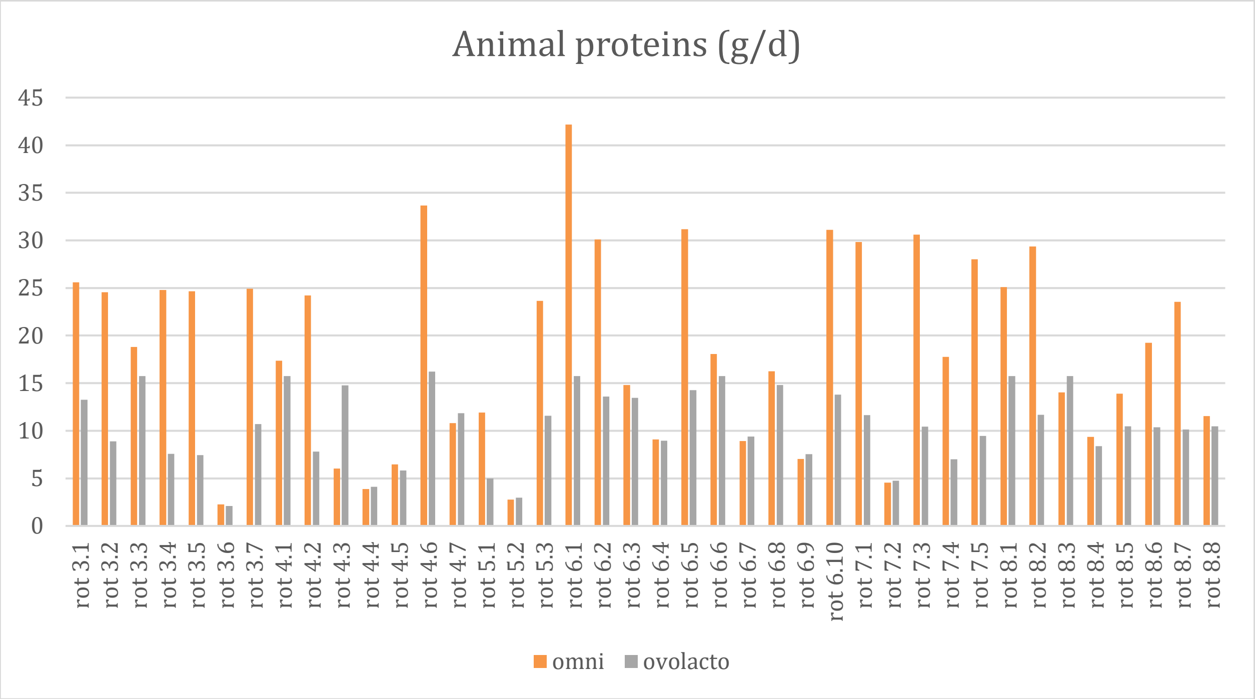


*Figure S.14. Amount of animal proteins (g/d) provided by the different rotations in the case of omnivorous (in orange) and ovo-lacto vegetarian (in grey) diets complying with the recommendations of the EAT-Lancet commission*

There was a clear correlation between the amount of Fe supplied and the production of eggs in all the rotations. Rotations supplying ovo-lacto vegetarian diets with eggs systematically had a higher animal iron production. Contents in animal protein were almost always higher in omnivorous diets, except for the rotations 4.3, 4.4, 4.7, 5.2, 6.7, 6.9, 7.2 and 8.3 producing more pork and lamb than beef as source of red meat. The share of animal proteins in the diets were also correlated with the herd sizes.
